# Evidence of Absence Regression: A Binomial N-Mixture Model for Estimating Fatalities at Wind Energy Facilities

**DOI:** 10.1101/2020.01.21.914754

**Authors:** Trent McDonald, Kimberly Bay, Jared Studyvin, Jesse Leckband, Amber Schorg, Jennifer McIvor

## Abstract

Estimating bird and bat fatalities caused by wind-turbine facilities is challenging when carcass counts are rare and produce counts that are either exactly zero or very near zero. The rarity of found carcasses is exacerbated when live members of a particular species are rare and when carcasses degrade quickly, are removed by scavengers, or are not detected by observers. With few observed carcass counts, common statistical methods like logistic, Poisson, or negative binomial regression are unreliable (statistically biased) and often fail to provide answers (i.e., fail to converge). Here, we propose a binomial N-mixture model that estimates fatality rates as well as the total number of carcass counts when these rates are expanded. Our model extends the ‘evidence of absence’ model (Huso et al., 2015; Dalthorp, Huso, and Dail, 2017) by relating carcass deposition rates to study covariates and by incorporating terms that naturally scale counts from facilities of different sizes. Our model, which we call Evidence of Absence Regression (EoAR), can estimate the total number of birds or bats killed at a single wind energy facility or a fleet of wind energy facilities based on covariate values. Furthermore, with accurate prior distributions the model’s results are extremely robust to sparse data and unobserved combinations of covariate values. In this paper, we describe the model, show its low bias and high precision via computer simulation, and apply it to bat carcass counts observed at 21 wind energy facilities in Iowa.

## 1 Introduction

The number of birds and bats killed by wind turbine blade strikes in North America and Europe each year varies wildly. A review of 116 studies from across the United States found between 0.27 and 11.02 small passerine deaths per megawatt of capacity per year (Erickson et al., 2014, Table 2). Given the wide range and sometimes low number of carcasses, particularly when partitioned into species groups or focused on endangered species, it is not surprising that the number of carcasses available to the analyst is often either zero or too few to make meaningful estimates. In addition, searchers find only a fraction of the carcasses at a site due to carcass decomposition, removal by scavengers, detection failure, or failure to search everywhere that carcasses lie and this only exacerbates the analysis difficulties inherent in low counts.

**Table 1:**
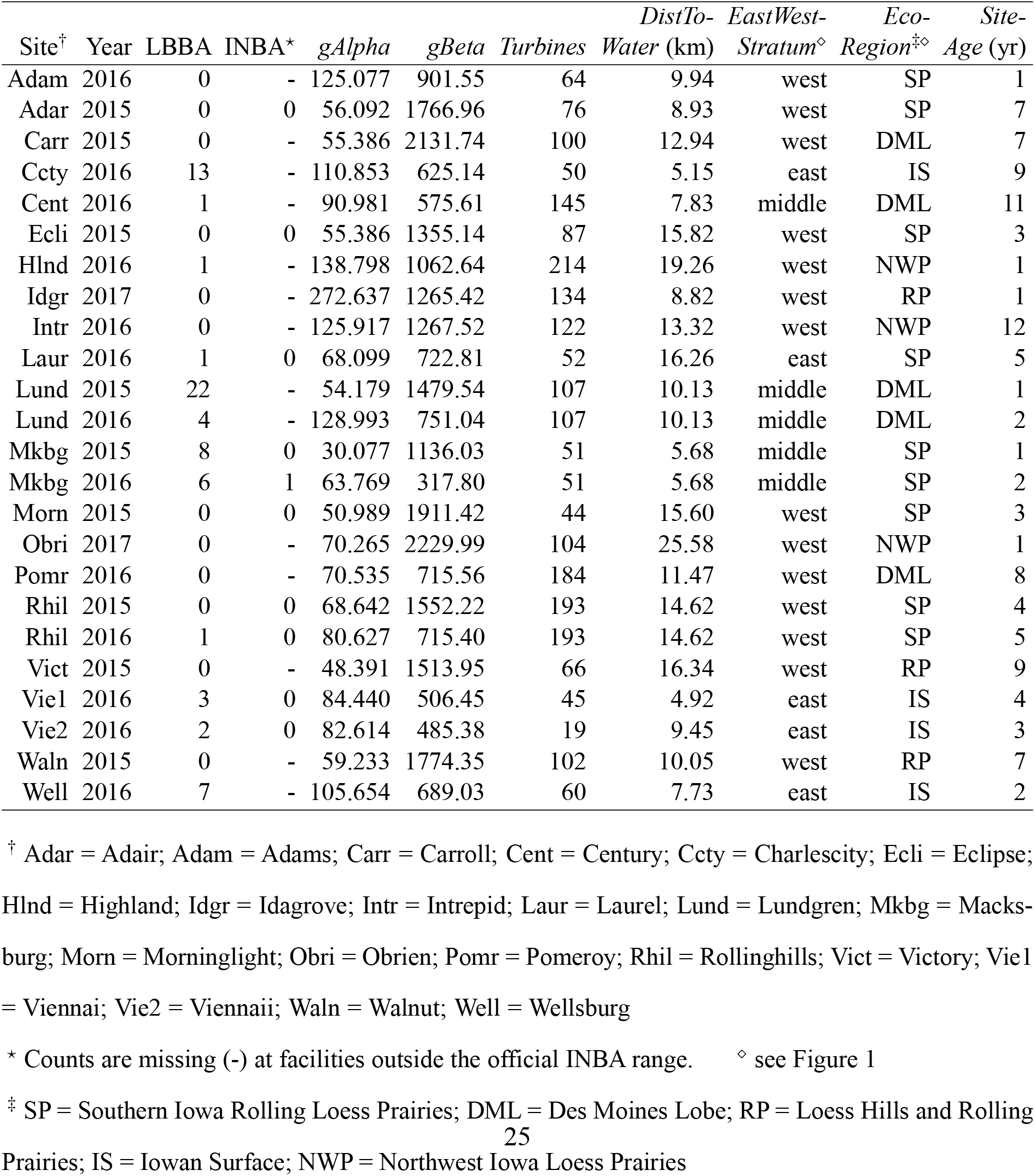
Wind facility monitoring data collected in Iowa (USA). **LBBA** and **INBA** columns contain, respectively, the number of little brown bat (*Myotis lucifugus*) and Indiana bat (*Myotis sodalis*) carcasses found at the site during all visits that year. *gAlpha* and *gBeta* are parameters of the Beta distribution for *g* estimated by R package `GenEst`. ***Turbines*** is number of wind turbines at the site. ***DistToWater*** is the average distance of turbines to nearest water (river, lake, or swamp) in kilometers. ***EastWestStratum*** is east-west stratum membership. ***EcoRegion*** is ecologic sub-region of the site. ***SiteAge*** is number of years the facility has been operating.

| Site<sup>†</sup> | Year | LBBA | INBA<sup>*</sup> | *gAlpha* | *gBeta* | Turbines | DistTo-Water (km) | EastWest-Stratum<sup>◇</sup> | Eco-Region<sup>‡◇</sup> | Site-Age (yr) |
| --- | --- | --- | --- | --- | --- | --- | --- | --- | --- | --- |
| Adam | 2016 | 0 | - | 125.077 | 901.55 | 64 | 9.94 | west | SP | 1 |
| Adar | 2015 | 0 | 0 | 56.092 | 1766.96 | 76 | 8.93 | west | SP | 7 |
| Carr | 2015 | 0 | - | 55.386 | 2131.74 | 100 | 12.94 | west | DML | 7 |
| Ccty | 2016 | 13 | - | 110.853 | 625.14 | 50 | 5.15 | east | IS | 9 |
| Cent | 2016 | 1 | - | 90.981 | 575.61 | 145 | 7.83 | middle | DML | 11 |
| Ecli | 2015 | 0 | 0 | 55.386 | 1355.14 | 87 | 15.82 | west | SP | 3 |
| Hlnd | 2016 | 1 | - | 138.798 | 1062.64 | 214 | 19.26 | west | NWP | 1 |
| Idgr | 2017 | 0 | - | 272.637 | 1265.42 | 134 | 8.82 | west | RP | 1 |
| Intr | 2016 | 0 | - | 125.917 | 1267.52 | 122 | 13.32 | west | NWP | 12 |
| Laur | 2016 | 1 | 0 | 68.099 | 722.81 | 52 | 16.26 | east | SP | 5 |
| Lund | 2015 | 22 | - | 54.179 | 1479.54 | 107 | 10.13 | middle | DML | 1 |
| Lund | 2016 | 4 | - | 128.993 | 751.04 | 107 | 10.13 | middle | DML | 2 |
| Mkgb | 2015 | 8 | 0 | 30.077 | 1136.03 | 51 | 5.68 | middle | SP | 1 |
| Mkgb | 2016 | 6 | 1 | 63.769 | 317.80 | 51 | 5.68 | middle | SP | 2 |
| Morn | 2015 | 0 | 0 | 50.989 | 1911.42 | 44 | 15.60 | west | SP | 3 |
| Obri | 2017 | 0 | - | 70.265 | 2229.99 | 104 | 25.58 | west | NWP | 1 |
| Pomr | 2016 | 0 | - | 70.535 | 715.56 | 184 | 11.47 | west | DML | 8 |
| Rhil | 2015 | 0 | 0 | 68.642 | 1552.22 | 193 | 14.62 | west | SP | 4 |
| Rhil | 2016 | 1 | 0 | 80.627 | 715.40 | 193 | 14.62 | west | SP | 5 |
| Vict | 2015 | 0 | - | 48.391 | 1513.95 | 66 | 16.34 | west | RP | 9 |
| Vie1 | 2016 | 3 | 0 | 84.440 | 506.45 | 45 | 4.92 | east | IS | 4 |
| Vie2 | 2016 | 2 | 0 | 82.614 | 485.38 | 19 | 9.45 | east | IS | 3 |
| Waln | 2015 | 0 | - | 59.233 | 1774.35 | 102 | 10.05 | west | RP | 7 |
| Well | 2016 | 7 | - | 105.654 | 689.03 | 60 | 7.73 | east | IS | 2 |

<sup>†</sup> Adar = Adair; Adam = Adams; Carr = Carroll; Cent = Century; Ccty = Charlescity; Ecli = Eclipse; Hlnd = Highland; Idgr = Idagrove; Intr = Intrepid; Laur = Laurel; Lund = Lundgren; Mkgb = Macksburg; Morn = Morninglight; Obri = Obrien; Pomr = Pomeroy; Rhil = Rollinghills; Vict = Victory; Vie1 = Viennai; Vie2 = Viennaii; Waln = Walnut; Well = Wellsburg
<sup>\*</sup> Counts are missing (-) at facilities outside the official INBA range. <sup>◇</sup> see Figure 1
<sup>‡</sup> SP = Southern Iowa Rolling Loess Prairies; DML = Des Moines Lobe; RP = Loess Hills and Rolling Prairies; IS = Iowan Surface; NWP = Northwest Iowa Loess Prairies

**Table 2:** All possible one, two, and three-variable models relating covariates (Table 1) to LBBA carcass counts after adjustment for facility-specific detection probabilities (*gAlpha* and *gBeta*) and facility size (*Turbines*). Fleet-wide mortality estimates and 90% credible interval limits are 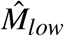, and 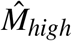, respectively. *df* is the number of coefficients in the model. Model 1 was chosen as the final model for reasons given in the text.

| Model | Structure | $df$ | DIC | $\hat{M}$ | $\hat{M}_{low}$ | $\hat{M}_{high}$ |
| --- | --- | --- | --- | --- | --- | --- |
| 1 | $\log(\lambda) \sim DistToWater + EastWestStratum + SiteAge$ | 5 | 107.36 | 672 | 548 | 826 |
| 2 | $\log(\lambda) \sim EastWestStratum + EcoRegion + SiteAge$ | 8 | 107.75 | 675 | 542 | 831 |
| 3 | $\log(\lambda) \sim EastWestStratum + SiteAge$ | 4 | 108.74 | 683 | 556 | 834 |
| 4 | $\log(\lambda) \sim DistToWater + EastWestStratum + EcoRegion$ | 8 | 117.53 | 637 | 520 | 786 |
| 5 | $\log(\lambda) \sim DistToWater + EcoRegion + SiteAge$ | 7 | 118.03 | 693 | 562 | 839 |
| 6 | $\log(\lambda) \sim EastWestStratum + EcoRegion$ | 7 | 121.95 | 624 | 508 | 756 |
| 7 | $\log(\lambda) \sim DistToWater + EastWestStratum$ | 4 | 126.26 | 629 | 510 | 757 |
| 8 | $\log(\lambda) \sim EastWestStratum$ | 3 | 127.55 | 633 | 520 | 766 |
| 9 | $\log(\lambda) \sim EcoRegion + SiteAge$ | 6 | 143.20 | 745 | 602 | 895 |
| 10 | $\log(\lambda) \sim DistToWater + EcoRegion$ | 6 | 156.52 | 665 | 547 | 807 |
| 11 | $\log(\lambda) \sim DistToWater + SiteAge$ | 3 | 162.07 | 632 | 522 | 764 |
| 12 | $\log(\lambda) \sim EcoRegion$ | 5 | 171.87 | 727 | 586 | 885 |
| 13 | $\log(\lambda) \sim DistToWater$ | 2 | 175.53 | 638 | 526 | 771 |
| 14 | $\log(\lambda) \sim SiteAge$ | 2 | 209.99 | 770 | 630 | 941 |
| 15 | $\log(\lambda) \sim 1$ | 1 | 222.03 | 793 | 653 | 956 |

There are several statistical techniques available for estimating fatality rates and total mortality when carcass counts are not small. Commonly applied techniques include the Horvitz-Thompson-type methods (Horvitz and Thompson, 1952; Särndal, Swensson, and Wretman, 1992; Lohr, 2010) of Shoenfeld (2004), Jain (2005), Korner-Nievergelt, Korner, et al. (2011), Huso (2011), and GenEst (Dalthorp, Madsen, et al., 2018; Dalthorp, Simonis, et al., 2020). These Horvitz-Thompson-type approaches generally estimate a single mean fatality rate (e.g., per turbine per year) and scale that mean by the number of turbines in order to estimate annual totals. When carcasses are common, techniques like normal-theory regression, generalized linear regression models (McCullagh and Nelder, 1989), negative binomial regression, and occupancy methods (MacKenzie, Nichols, Hines, et al., 2003; MacKenzie, Nichols, Sutton, et al., 2005; Pavlacky et al., 2012) can be used to model fatality rates as a function of covariates and hence account for variation due to measurable factors. Many of these techniques also account for variation in detection probabilities, either by including scaling terms (e.g., number of turbines, number of searches, length of searches) or by including detection as an explicit parameter.

But, when carcasses are rare, estimation of a mean fatality rate and its variance via regression is challenging, and all the regression models cited in the previous paragraph encounter difficulties. In particular, the increased prevalence of zeros causes model instability due to unobserved or low counts in certain covariate combinations. Unobserved covariate combinations cause what is known as complete or quasi-complete separation (Albert and Anderson, 1984) and is a problem because the empirical rate for unobserved combinations is exactly zero yet the modeled mean rate must be greater than zero. In other words, zeros or low counts in certain covariate combinations reduces or eliminates the amount of information available to estimate one or more coefficients in the model.

Bias is also a problem for many regression techniques that are applied to rare event data. Low variation near zero is known to cause a sharp underestimate of rare event probabilities (King and Zeng, 2001) which in turn leads to biased coefficient estimates. To combat this bias, analysts use weighted logistic regression (King and Zeng, 2001; Maalouf and Trafalis, 2011) and modifications of machine learning algorithms (Maratea, Petrosino, and Manzo, 2014). Both of these techniques use separate estimates of rare event probabilities to adjust and stabilize coefficient estimates, but still at least one success is required in all covariate combinations. Many times, separate estimates of rare event probabilities are not available and researchers must postulate a level without actually observing any events. Despite modifications that combat bias, nothing resolves logistic regression’s inability to estimate rates when zero successes are observed in one or more combinations of factor levels.

To combat problems associated with rarity when monitoring wind energy facilities after construction, Huso et al. (2015) introduced the probability of detection (here, denoted *g*) into a binomial N-mixture model (Kéry and Royle, 2016, Ch. 7) and used this model to estimate total fatalities. The key feature of Huso et al.’s (2015) model is that detection probabilities are estimated from the study design (e.g., survey frequency, carcass persistence, detection probability, etc.) and not from observed carcass counts in the binomial portion of the model. This feature allows the model to separate information on totals from information on detection probability and hence avoid the identifiability issues that plague some binomial N-mixture models (Barker et al., 2018; Kéry, 2018).

Huso et al.’s (2015) method is known as the Evidence of Absence (EoA) approach (see also Dalthorp, Huso, Dail, and Kenyon, 2014; Dalthorp, Huso, and Dail, 2017) and represents a major step forward in analysis of fatalities at wind facilities. Current parameterizations of the EoA mixture model focus on estimating the total number of fatalities that occurred during a particular year. Current EoA models do not provide estimates of covariate relationships between study factors and fatality rates, and accounting for differences among facility sizes is accomplished after separate estimation. The utility of EoA would be greatly enhanced if it estimated covariate relationships and if it included terms that allowed variable wind farm sizes in a single model run.

In this paper, we extend EoA to allow estimation of covariate relationships and we show that including the logarithm of facility size (e.g., number of turbines, mega-watts, footprint) as a term without an estimated coefficient naturally scales the model’s rate estimates to accommodate different facility sizes. Our main extension is inclusion of an additional level of hierarchy, relative to EoA, in our binomial N-mixture model that parameterizes a log-linear relationship between study covariates and sample-unit fatality rates. This extension allows factor and continuous effects to act upon rate estimates, thereby reducing unmodeled heterogeneity. While not necessary, we also include scaling factors, commonly called offset terms (McCullagh and Nelder, 1989, Chapter 6), in the log-linear model to scale rate estimates to natural units (e.g., per turbine, per hectare, per year, etc.) based on the monitoring protocol for each wind farm. It is then possible to derive estimates of total fatalities across space or time by simple scaling operations. If zero mortalities are observed, it is possible to inform the prior distribution of our model’s rate parameter using previous estimates or separate information and thereby ameliorate or eliminate the separation issues that plague other techniques. If zero carcasses are observed and it is not possible to inform the rate parameter’s prior, the method still provides reasonable fatality estimates because the information provided by detection probabilities (i.e., *g*) is being considered and this stabilizes the estimation process greatly. While our focus is primarily on bird and bat fatality rates, the model is useful in any situation where detection probabilities can be quantified. Our model is particularly useful when carcass counts are appreciable (e.g., ≥ 1 in most covariate groups) and covariate relationships are sought. We call our model Evidence of Absence Regression, or EoAR, to reflect the fact that this model is essentially EoA with a regression relationship tacked on.

In the remainder of this article, we define the EoAR hierarchical model and provide examples of the model’s application to bat carcass counts at wind energy facilities in Iowa. We also describe several sets of statistical simulations that investigate EoAR’s statistical properties.

## 2 Methods

### 2.1 EoAR Model Definition

We define a site to be the temporal and spatial units over which observed counts are summarized. For example, studies of wind energy facilities typically summarize carcass counts by facility and year (i.e., site = annual facility). Alternatively, carcass counts could be summarized by turbine and field season (i.e., site = seasonal turbine) or by turbine over all years of the study (i.e., site = turbine).

We assume *m* sites are included in the study and that each site is searched a variable number of times over a variable length of time. During each search, field personnel record *c*_*i j*_, the number of carcasses detected during search *j* of site *i*. The count summary we analyze is the total number of observed events at site i summed over searches that occurred during the monitoring period, *C*_*i*_ = ∑ _*j*_ *c*_*i j*_. To accommodate sites of different sizes and monitoring periods of differing lengths, we define *A*_*i*_ to be the number of natural units (e.g., days, years, turbines, facilities, facility-years, etc.) over which past data was collected at site *i*. Typically, *C*_*i*_ is the total number of carcasses observed at facility *i* during a particular year and *A*_*i*_ is the number of turbines monitored at facility *i*.

The probability of detection (*g*) is a key parameter, both here and elsewhere (Huso et al., 2015; Dalthorp, Huso, Dail, and Kenyon, 2014; Dalthorp, Huso, and Dail, 2017; Dalthorp, Madsen, et al., 2018), and its estimation for wind facility monitoring studies is too complicated to detail here. Wind facility carcass search detection probabilities are complicated because they depend on the timing of seasonal carcass arrivals, start date of searches, end date of searches, search interval, number of searched turbines, searcher efficiency (detection given presence), carcass removal rates, and the proportion of the carcasses that fall in the searched area. Here, we rely on established formulas implemented in the `GenEst` R package (Dalthorp, Simonis, et al., 2020) to compute detection probabilities and their distribution given study design elements (Dalthorp and Huso, 2015; Reyes et al., 2016).

Whether using `GenEst` or not, we assume site-specific *g*_*i*_ are available. We recommend using site-specific *g*_*i*_ when available, but acknowledge there may be data poor situations in which it is reasonable to assume certain *g*_*i*_ are random variables arising from a single distribution. If detection probabilities are pooled in this way, we recommend assuming the pooled *g*’s arise from an aggregate beta distribution and that practitioners compute a weighted average mean *g* for the aggregate distribution as,

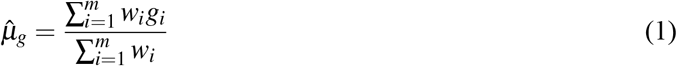

where weights *w*_*i*_ should theoretically be proportional to the total number of mortalities at site *i*. In practice, total mortality at all sites is often unknown and it is common to set *w*_*i*_ = *A*_*i*_ because *A*_*i*_ is often theoretically proportional to the largest risk factor, e.g., size. We also recommend a weighted standard deviation for the aggregate *g* distribution computed as,

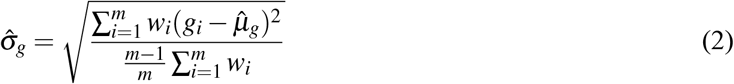

Given 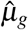 and 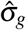, we estimate the *α* and *Ψ* parameters of *g*’s aggregate Beta distribution using method of moments,

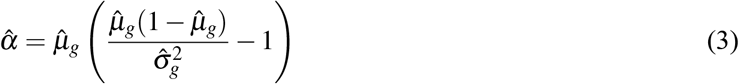

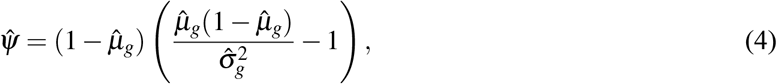

which assumes 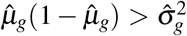. Estimation of *α* and *Ψ* using maximum likelihood and individual *g*_*i*_’s is also possible but is more complicated. If using method of moments and 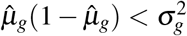, we evaluate the histogram of *g*_*i*_ and either reduce 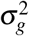 slightly or discard the Beta distribution in favor of a truncated Gamma or a smoothed histogram density estimate (Silverman, 1998).

Our interest lies primarily in the site-specific rate, *λ*_*i*_, defined as the number of events per units of *A*_*i*_. Again, *λ*_*i*_ in wind facility monitoring studies typically estimates fatalities per turbine per year, but could be fatalities per facility-year, per turbine over multiple years, per turbine per month, etc. The most useful basis for *λ*_*i*_ depends largely on the utility and availability of appropriate covariates.

When *g* < 1, we do not know the true number of mortalities, only that it is greater than or equal to *C*_*i*_. Associated with *λ*_*i*_ in this case is the true number of mortalities at site *i* over *A*_*i*_ units, which we define to be *M*_*i*_ (*M*_*i*_ ≥ *C*_*i*_). To estimate *M*_*i*_, we employ a Bayesian approach which views both *λ*_*i*_ and *M*_*i*_ as random variables, and in particular assumes that *M*_*i*_ follows a Poisson distribution with mean *λ*_*i*_*A*_*i*_. Given values of *M*_*i*_, the total number of past fatalities over all sites is 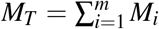. Assuming independence of carcass counts among sites, *M*_*T*_ is Poisson with mean 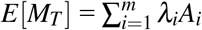.

The key feature of EoAR is the regression relationship between *λ*_*i*_ and exogenous site-specific covariates, like season, habitat, year, distance, etc. By estimating such a relationship, we allow heterogeneity in rates across sites and predictions of future fatalities are correspondingly improved. The hierarchical binomial N-mixture EoAR model is,

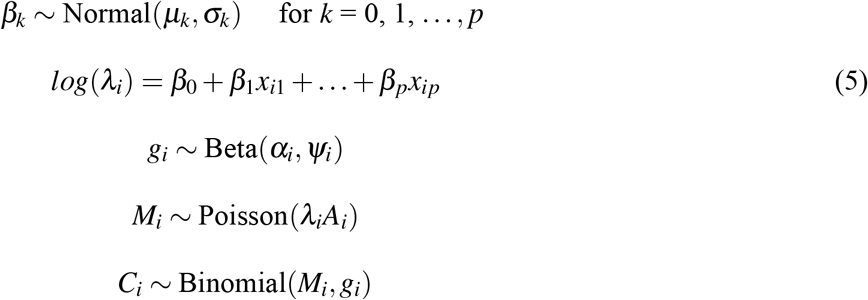

In this model, the *β*_*k*_’s are coefficients to be estimated, while *α*_*i*_ and *Ψ*_*i*_ are constants. To illustrate the model visually, Appendix S1: Figure S1 contains a directed acyclic graph depicting the structure of EoAR, drawn for Bayesian analysis along the lines suggested by Kruschke (2011). We choose to estimate coefficients in Equation 5 using MCMC sampling implemented in JAGS (Plummer, 2003), but other estimation techniques are possible (e.g., STAN or direct evaluation). We generally use the median of a parameter’s posterior distribution as that parameter’s point estimate and quantiles of the posterior as the parameter’s credible interval, but other summaries of the posteriors are possible (e.g., modes, means, and highest-density intervals). An R package for estimating EoAR models using formula-based specification is available from the authors and via links provided in Appendix S1.

If we have prior estimates or independent information about the location and width of the distribution for certain *β* coefficients, it can be incorporated into the EoAR model by setting *µ*_*k*_ and *σ*_*k*_ accordingly. A special case arises when only an intercept is present in Equation 5 and independent information exists on fatality rate. Suppose we obtain independent information regarding the mean rate, 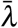and its variation over all *m* sites. For example, perhaps we obtain an estimate of total mortalities, 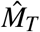, and its standard deviation, 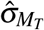, that can appropriately be used as prior information. A reasonable prior estimate of 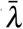 is then 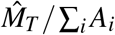, and a prior estimate of 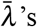 standard error is 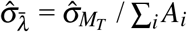. Assuming 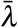 follows a log-normal distribution, the intercept parameter in Equation 5 will follow a normal distribution, and we can set the prior mean and standard deviation of the intercept using the following equations,

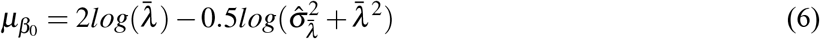

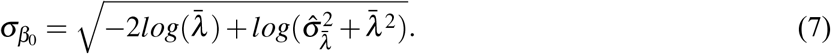

When prior information on coefficients is not available, we choose to use vague prior distributions for the *β*_*k*_’s. To implement vague priors, we set *µ*_*k*_=0 and *σ*_*k*_ equal to 100 times the upper limit of a (Wald) 95% confidence limit for *β*_*k*_ obtained from a Poisson regression of *C*_*i*_/*g*_*i*_ on *x*_*i*1_ through *x*_*ip*_. Specifically, our vague priors set *σ*_*k*_ = 100(|*b*_*k*_ + 2*sd*_*k*_|) where *b*_*k*_ and *sd*_*k*_ are the estimated coefficient and standard error for *β*_*k*_ in a Poisson regression containing covariates and offset term *log*(*g*_*i*_) + *log*(*A*_*i*_). If the Poisson regression does not converge due to incomplete separation, we remove effects without variation in *C*_*i*_ until the regression converges and set prior standard deviations for the coefficients of removed effects equal to 100.

### 2.2 Simulations

To investigate bias and precision of EoAR’s *λ* and *M* estimators, we ran three sets of simulations, one set without covariates, one set with a simple factor covariate, and a final set with variable facility sizes.During the first set of simulations, we set the true *β*_0_ = 0 to simulate a true fatality rate of *λ* = 1. We set the number of sites, *m*, to 20 and 50 and the true mean detection probability to *µ*_*g*_ = 0.1 and 0.3. For both levels of mean detection probability, the simulation assumed a variance of 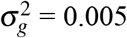 and computed *α* and *Ψ* using Equations 3 and 4. At all sites (*i* = 1, 2,…, *m*), the simulation generated actual fatalities using a Poisson distribution, i.e.,

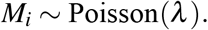

Finally, the simulation generated observed carcass counts using the binomial distribution, i.e.,

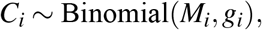

where *g*_*i*_ ~ Beta(*α, Ψ*). Given *C*_*i*_, *α*, and *Ψ*, we estimated *λ, M*_*i*_ and *M*_*T*_ = ∑_*i*_ *M*_*i*_ using the EoAR model outlined in Section 2.1. We computed point estimates as the median of each posterior distribution and reported histograms of the rate parameter’s point estimates, 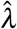, and fatality estimation errors, 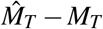, over 500 iterations of the simulation. We estimated bias of both 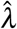 and 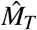 as the numerical average of each estimator over iterations minus the true value.

Our second set of simulations was similar to the first except that we added a single discrete factor with two levels to the equation for true *λ*. This simulation investigated whether EoAR could recover rate information when carcass counts were rare at some sites and common at others. We set the factor covariate, *x*_*i*1_, to 0 for half the sites in the simulation (i.e., *x*_*i*1_ = 0 for *i* = 1, 2,…, *m*/2) and 1 for the other half. To simulate high and low rates among sites, we set the true *β*_0_ = log(10) = 2.3026 and *β*_1_ = −log(10) = −2.3026. These *β* values translated into rates of *λ*_*i*_ = 10 for the first half of the simulated sites and *λ*_*i*_ = 1 for the second half. We set the remaining parameters to make the second simulations comparable to the first. That is, we set *m* to 20 and 50, *µ*_*g*_ = 0.1 and 0.3, simulated *M*_*i*_ ~ Poisson(*λ*_*i*_), and *C*_*i*_ ~ Binomial(*M*_*i*_, *µ*_*g*_).

Our third set of simulations extended the second to investigate whether EoAR could recover information on two *λ* values in the face of extreme disparities in carcass counts. All runs of our third simulation assumed *m* = 2 sites, one with *λ*_*i*_ = 1 that contained 40 turbines and one with *λ*_*i*_ = 10 that contained 10 turbines. During one run of this simulation, site 1 had an average *g* value of 0.1 while the second site had an average *g* value of 0.9. That equated to an average carcass count of 4 at site 1 (*A*_1_*λ*_1_*g*_1_ = 40(1)(0.1)) and 90 at site 2 (*A*_2_*λ*_2_*g*_2_ = 10(10)(0.9)). During a second run, we assigned site 1 an average *g* value of 0.9 and assigned site 2 an average *g* value of 0.1. The second run’s parameters equated to an average carcass count of 36 at site 1 (*A*_1_*λ*_1_*g*_1_ = 40(1)(0.9)) and 10 at site 2 (*A*_2_*λ*_2_*g*_2_ = 10(10)(0.1)). Our main question during these runs was whether EoAR could accurately estimate fatality rates (i.e., *λ*_1_ = 1 and *λ*_2_ = 10) despite drastic differences in the number of carcasses observed at each site.

R code to carry out the simulations is included in supplemental files DataS1:EoARSimulation1.R, DataS1:EoAR-Simulation2.R, and DataS1:EoARSimulation3.R

## 3 Bat Fatality Monitoring in Iowa

To illustrate EoAR numerically, we apply it to Indiana bat (INBA) (*Myotis sodalis*) and little brown bat (LBBA) (*Myotis lucifugus*) carcass data collected during 2015, 2016, and 2017 at twenty-one operating wind energy facilities located in Iowa (Figure 1). We choose INBA and LBBA for these examples, but species of the animal of interest has no bearing on the analysis. EoAR can be run on any species or collection of species. We conducted these studies in part because INBA are listed as endangered under the Endangered Species Act and LBBA are considered rare and of concern. LBBA are considered to occur statewide in Iowa, whereas INBA are considered by regulators to occur only in the southeast quarter (approximately) of the state.

**Figure 1:**
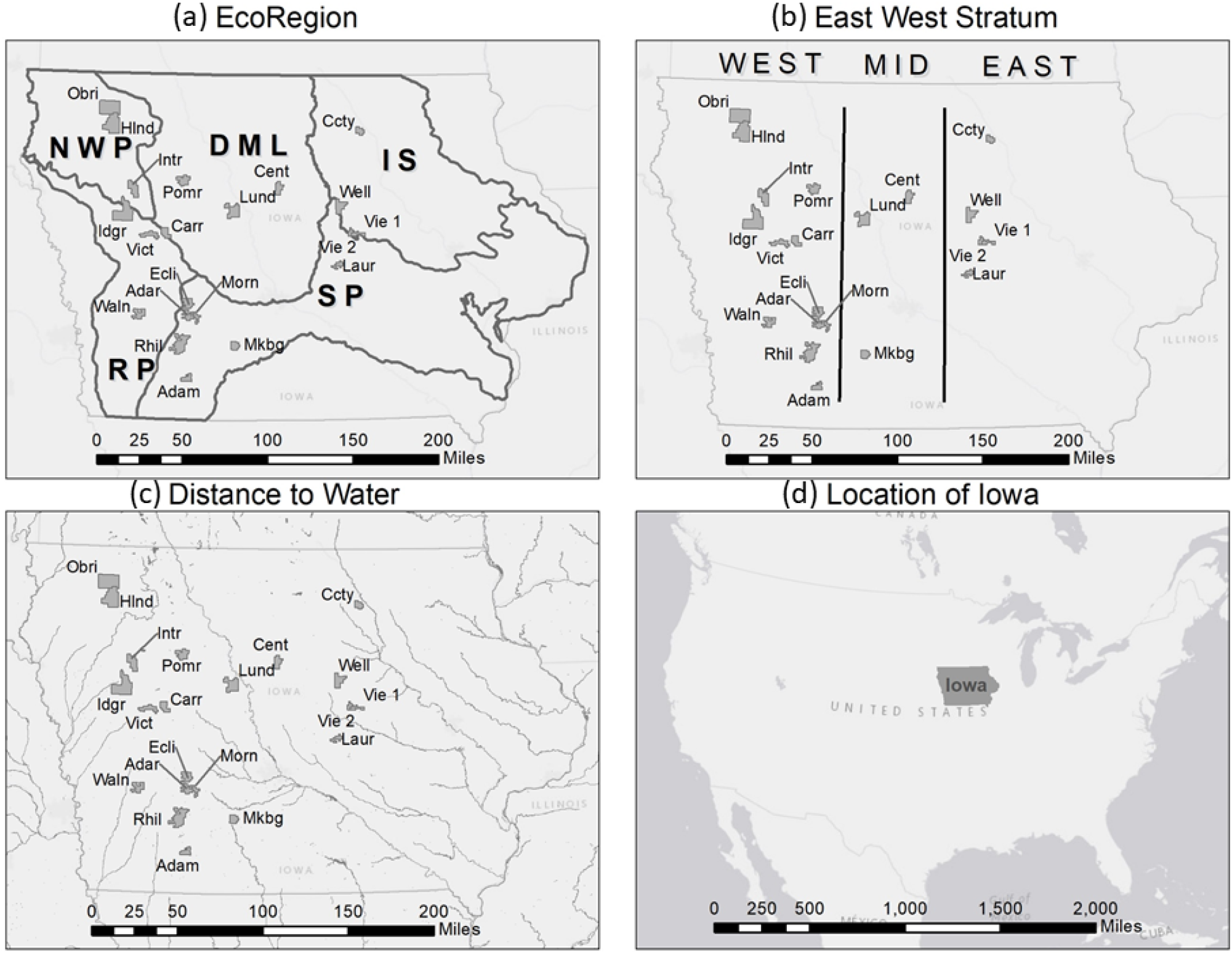
Geographic covariates used in Example 1. (a) ecological sub-regions, (b) east-west stratum covariate boundaries, (c) rivers and lakes in Iowa used to derive the distance to water covariate, (d) the location of Iowa within the United States. Grey polygons are wind facilities. Facility abbreviations and corresponding covariate values appear in Table 1.

Field data collection occurred from 15 March through 15 November in 2015 through 2017, and not all facilities were monitored every year. During data collection, field personnel searched for bat carcasses every 7 days (2015) or 3 days (2016 and 2017) beneath turbines on plots of varying size and shape. In 2015, personnel walked the perimeter of turbine pads that encompassed approximately 10m of gravel surrounding the turbine base, and along the access road to a distance of 100m from the turbine. In 2016 and 2017, contractors mowed square plots centered on the turbine at a random sample of 20% of the turbines. The sizes of mowed plots were 60m X 60m, 100m X 100m, or 200m X 200m. We term the mowed plots ‘full’ plots and technicians searched them by walking straight transects that traversed their width and that were spaced 10m apart. At the other 80% of turbines in 2016 and 2017, technicians walked the perimeter of the turbine pad and along the access road out to a distance of 100m from the turbine. We collected data at nine facilities in 2015, thirteen facilities in 2016, and two facilities in 2017. We visited three facilities in both 2015 and 2016 (Table 1).

In the remainder of this section, we describe two example applications that demonstrate EoAR’s flexibility. In the first, we relate annual counts of LBBA to study covariates and identify factors affecting fatalities per turbine per year. The LBBA model contained covariates because search efforts produced carcasses of this species with enough frequency to support inclusion of covariates. In the second example, we analyze INBA counts, which were less frequently found than LBBA, to illustrate estimation of the model without covariates. In the second example, we fit an intercept-only model with and without informed prior distributions to illustrate the consequences of informing the intercept’s distribution.

### 3.1 Example 1: LBBA

The covariates we considered in this example included ecological sub-region of the facility (Figure 1a), an east-west grouping of facilities we termed ‘stratum’ (Figure 1b), distance to the nearest river greater than class 4 (Figure 1c), and facility age (Table 1, column 11). Ecological sub-regions generally contain different vegetation communities and hence potentially different bat prey (insect) communities. The east-west stratum generally represented a transition from hardwood dominated woodlands in the east to more open grassland species in the west and hence potentially different communities of bats. Rivers are generally heavily wooded in Iowa and generally represent both bat roosting and foraging habitat. We included facility age, quantified as the number of years post-construction, because bat mortalities observed at other facilities have occasionally decreased after one or two years of operations.

We computed facility and year-specific detection probabilities (*g*_*i*_) using functions in `GenEst` (Dalthorp, Simonis, et al., 2020). We based estimates of detection probabilities (columns *gAlpha* and *gBeta*, Table 1) on visit timing, visit frequency, searcher efficiency, carcass persistence, and the proportion of the carcass’ spatial distribution that was searched. We derived both searcher efficiency and carcass persistence from field trials that involved placing carcasses at known locations and assessing detection (efficiency) and persistence. We estimated the searched proportion of the carcass distribution by selecting the best parametric curve fitted to the histogram of distance-to-turbine using a truncated weighted maximum likelihood approach implemented in the windAC R package (Studyvin, Rabie, and Riser-Espinoza, 2020).

For this example, we fitted all possible combinations of study covariates up to a maximum of three variables in the log-linear model for *λ* (Equation 5). All combinations of covariates *DistToWater, EastWestStratum, EcoRegion*, and *SiteAge* with three or fewer variables yielded 15 possible models (i.e., intercept only, intercept + *DistToWater*, intercept + *EastWestStratum*, …, intercept + *EastWestStratum* + *EcoRegion* + *SiteAge*). We assessed the fit of each model using the deviance information criterion (DIC) (Gelman et al., 2004; Spiegelhalter et al., 2014) and considered models within approximately 2 DIC units to be equivalent. We included the logarithm of the number of turbines at each facility (*Turbines*, Table 1) as an offset in all models.

### 3.2 Example 2: INBA

A single INBA carcass was found during monitoring at the eight facilities in the official INBA range, and this low number of carcasses could not support inclusion of a covariate in the EoAR model without drastically informing all coefficient priors. To illustrate the flexibility of EoAR, we fit both an uninformed and informed EoAR model under three scenarios for the distributions of *g*.

The first *g* scenario we fitted assumed that all facility and year-specific detection probabilities were simply random observations from a single underlying beta distribution. The second *g* scenario assumed that facility-specific detection probabilities were random observations from year-specific beta distributions. We took this second approach because crews surveyed roads and pads in 2015 but full plots and roads and pads in 2016. We used Equations 1 and 2 to compute the mean and standard deviation of each beta distribution (*w*_*i*_ = *A*_*i*_ = number of turbines). We then used Equations 3 and 4 to compute the *α* and *Ψ* parameters of each distribution. Lastly, we assumed that all facility and year-specific detection probabilities were separate independent random variables from beta distributions centered on the *g*_*i*_ point estimates. We feel the second *g* scenario is the most likely for this data set due to unexplained random variation in searcher efficiency at different sites within years, but our purpose is simply to illustrate the flexibility of EoAR. We also recognize that un-modeled heterogeneity in *g* can induce underestimation, and we caution readers to think carefully about the consequences of combining *g*’s across sites. If over-estimates can be tolerated more easily than under-estimates, it is prudent to use separate *g*_*i*_ for all sites.

Under each *g* scenario, we estimated the mean and upper bound of INBA fatalities from an intercept-only EoAR model with both vague and informed priors. A previous analysis of INBA fatalities in Iowa found 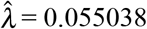 INBA fatalities per turbine per year with a standard deviation of 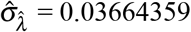. Using these values in Equations 6 and 7, we obtained prior estimates of 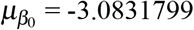 and 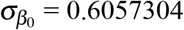.

Again, our aim in this example is to show EoAR’s flexibility. The incorporation of prior information is one of the advantages of Bayesian analyses over frequentist analyses, but inappropriate prior information can lead to biased results and erroneous conclusions. A full discussion of prior information, when it is appropriate, and the consequences of using inappropriate priors is beyond the scope of this paper. We refer the reader to discussions on selecting appropriate prior information in Lemoine (2019), Simpson et al. (2017), and Korner-Nievergelt, Roth, et al. (2015). Ideally, practitioners have resources to test prior information and its effects via simulation of realistic but challenging scenarios and can justify prior information on ecological or study design arguments.

## 4 Results

### 4.1 Example 1: LBBA

The DIC statistic ranked three of the fifteen fitted models as functionally equivalent (Table 2). DIC differed by only 1.38 between the top three models, while DIC differed by 8.8 units between the third and fourth-ranked model. The top three models contained both *EastWestStratum* and *SiteAge*, while the top model contained *DistToWater* and the second model contained *EcoRegion*. We felt the top ranked model that included *DistToWater* was reasonable, biologically plausible, and easily applied to study areas outside Iowa if deemed appropriate. For these reasons, we established the top-ranked model as our final model and illustrate the types of inferences that EoAR makes possible.

Coefficients in the model (Table 3) estimated a 7.8% drop in mortalities per turbine for every one kilometer increase in the average distance of a facility’s turbines from the nearest water (0.075 = 1 - exp(−0.078), 90% CI = 0% to 15%). To illustrate one use of the model, consider the case of an electric utility company contemplating installation of two new wind-energy facilities. Suppose the utility is considering installation of Facility A in the eastern stratum 5 kilometers from water and installation of Facility B in the same stratum 15 kilometers from water. The utility plans to build both facilities in the same year so site ages will be identical. The final EoAR model predicts that LBBA fatalities per turbine per year at Facility B will be on average 54.17% lower than at Facility A due to the difference in distance to water (0.5417 = 1 - exp(−0.078(10))). In other words, whatever Facility A’s fatality rate is, this model predicts that Facility B’s fatality rate will be *exp*(−0.078(10)) = 0.4583 times Facility A’s. The utility could also construct confidence and prediction bounds for the mortality rate and totals at Facility A, Facility B, or the difference in rates and totals using the EoAR model.

**Table 3:** Coefficients in the top-ranked (by DIC) EoAR model in Table 2. The model relates LBBA fatality rates to *DistToWater*, the *EastWestStratum* factor, and *SiteAge*. CI_5_ and CI_9_5 are endpoints of 90% credible intervals.

| Effect | Estimate | SD | $CI_5$ | $CI_{95}$ |
| --- | --- | --- | --- | --- |
| (Intercept) | 1.2755 | 0.4883 | 0.4960 | 2.0580 |
| *DistToWater* | -0.0780 | 0.0524 | -0.1673 | 0.0027 |
| *EastWestStratum.middle* | -0.0490 | 0.2774 | -0.5077 | 0.4067 |
| *EastWestStratum.west* | -4.0101 | 0.8841 | -5.6572 | -2.8436 |
| *SiteAge* | -0.1844 | 0.0458 | -0.2626 | -0.1124 |

We found only 2 LBBA carcasses in the western stratum, but 41 and 26 carcasses in the middle and eastern stratum, respectively. After factoring in facility-specific detection probabilities and that the middle stratum contained more turbines, the final model estimated that annual LBBA fatalities rates in the middle and eastern stratum were not statistically different (5% decrease from east to middle, 90% CI = −50% to 40%), but that rates decreased by 98% (90% CI = 96% to 99%) in the western stratum relative to the middle stratum.

The final model also estimated that fatalities of LBBA decreased as facilities aged. The final model estimated that LBBA fatality rates decreased by an average of 16.8% for every year the facility operated (90% CI = 10.6% to 23.1%). To illustrate, fatalities for a facility in the eastern stratum located 10 km from the nearest water was estimated at 1.3647 individuals per turbine per year during the facility’s first year of operation. After 10 years, the estimate of average annual fatalities at this same facility drops to 0.2595 individuals per turbine. After 30 years, the model estimates an average of 0.0065 annual fatalities per turbine at this facility.

### 4.2 Example 2: INBA

Applying Equations 1 through 4, the *α* and *Ψ* parameters of the single aggregate beta distribution were 541.61 and 6169.60, respectively (mean *g*_*i*_ = 0.081, *σ*_*g*_ = 0.0033), over all facility-year combinations. The *α* and *Ψ* parameters of the 2015 beta distribution for *g* were 254.25 and 7489.61, respectively (mean *g*_*i*_ = 0.033, *σ*_*g*_ = 0.0020). The *α* and *Ψ* parameters of the 2016 beta distribution were 359.01 and 2433.25, respectively (mean *g*_*i*_ = 0.129, *σ*_*g*_ = 0.0063).

Estimates of total mortality and fatality rates of INBA under vague and informed priors, as well as the three scenarios for *g*, appear in Table 4. We note that estimates from the informed model were almost 3 times higher than those from the uninformed model, and 90% credible intervals were approximately the same width. A naive and simple estimate of INBA deaths per turbine in this example is the number of carcasses divided by average *g* divided by number of turbines, or 1/0.081/811 = 0.0152. This naive estimate is closest to 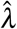 from the uninformed models with facility and year-specific *g* values. While it is difficult to ascertain MCMC simulation error in this example, this supports the notion that grouping *g* values can induce downward bias in rate estimates. As expected, the 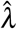 values from the informed models (~ 0.032) were approximately half-way between the prior estimate (0.0550) and the naive estimate (0.0152).

**Table 4:** Examples of estimates of total mortality 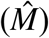 and fatality rate 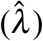 of INBA under vague and informed priors as well as three scenarios for the distributions of detection probabilities (*g*).

| Prior for<br>$\beta_0$ | Number of<br>$g$ groups | $\hat{M}$ | 90% CI for $M$ | | $\hat{\lambda}$ | 90% CI for $\lambda$ | |
| --- | --- | --- | --- | --- | --- | --- | --- |
|  |  |  | Lower | Upper |  | Lower | Upper |
| vague | 1 | 9 | 1 | 35 | 0.01048 | 0.00068 | 0.04483 |
| vague | 2 | 9 | 1 | 39 | 0.01145 | 0.00093 | 0.04843 |
| vague | 10 | 10 | 1 | 42 | 0.01251 | 0.00089 | 0.05151 |
| informed | 1 | 24 | 10 | 49 | 0.03092 | 0.01408 | 0.06212 |
| informed | 2 | 24 | 11 | 52 | 0.03132 | 0.01419 | 0.06555 |
| informed | 10 | 25 | 11 | 52 | 0.03258 | 0.01442 | 0.06719 |

Under all *g* scenarios, the standard deviation of the intercept parameter dropped by approximately 50% when we informed the prior. The corresponding standard deviation of 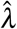 did not decrease by this amount because *λ* was close to zero and transformation through the exponential link mitigated the precision gained by informing the intercept’s prior. In other words, informed priors affect precision on the log-link scale, but only lead to corresponding changes in the precision of 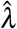 if rates are well above zero.

### 4.3 Simulation Results

Simulations containing a single low event rate produced uni-modal distributions for both 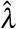 and estimation errors 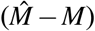 that surround the total number (Figure 2). The distributions of 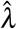 were discrete for *M* = 20 or *µ*_*g*_ = 0.1, but otherwise showed decreasing bias and increasing precision as either *M* or *µ*_*g*_ increased (Figure 2, Table 5). The distribution of estimation errors, 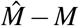, was centered on 0 and only slightly skewed right (slight underestimation) for the low level of *M* combined with the low level of *µ*_*g*_. The estimated coverage of credible intervals for both *λ* and *M* were slightly below (generally ≤ 3%) the nominal level of 90% and approached the nominal level as *µ*_*g*_ or true *M* increased (Table 5).

**Table 5:** Percent bias and coverage of 90% credible intervals for 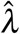 and 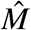 computed during both sets of simulations. *λ* is the average number of fatalities per site. *m*_*i*_ is the number of sites. *g*_*i*_ is the mean probability of detecting a fatality. Simulation 3 contained one site with *A*_*L*_ = 40 turbines and one site with *A*_*H*_ = 10 turbines, and each site had the corresponding *g* specified in the fourth column. The true number of fatalities was Poisson(*A*_*i*_*λ*_*i*_) at each site and averaged ∑ *m*_*i*_*λ*_*i*_ when summed over sites.

| Simulation | $m_i$ | $A_i$ | $g_i$ | $\hat{\lambda}_L = 1$ | | $\hat{\lambda}_H = 10$ | | $\hat{M}$ | |
| --- | --- | --- | --- | --- | --- | --- | --- | --- | --- |
|  |  |  |  | %Bias | %Cover | %Bias | %Cover | %Bias | %Cover |
| 1 | 20 | 1 | 0.1 | -16.47 | 87.00 | - | - | -16.13 | 86.80 |
| 1 | 20 | 1 | 0.3 | -7.81 | 87.00 | - | - | -5.76 | 87.60 |
| 1 | 50 | 1 | 0.1 | -5.26 | 84.40 | - | - | -4.73 | 88.60 |
| 1 | 50 | 1 | 0.3 | -1.99 | 87.20 | - | - | -2.13 | 89.80 |
| 2 | 20 | 1 | 0.1 | -16.45 | 62.40 | -2.13 | 97.00 | -1.24 | 95.80 |
| 2 | 20 | 1 | 0.3 | -11.12 | 85.40 | -1.34 | 92.60 | -0.76 | 93.80 |
| 2 | 50 | 1 | 0.1 | -7.46 | 89.00 | -1.12 | 94.40 | -0.92 | 94.60 |
| 2 | 50 | 1 | 0.3 | -7.06 | 90.00 | -0.26 | 92.60 | -0.46 | 93.60 |
| 3 | 1,1 | 40,10 | 0.1,0.9 | 9.44 | 97.60 | -0.52 | 93.80 | 3.82 | 97.80 |
| 3 | 1,1 | 40,10 | 0.9,0.1 | 0.83 | 92.40 | 18.66 | 99.80 | 13.64 | 99.80 |

**Figure 2:**
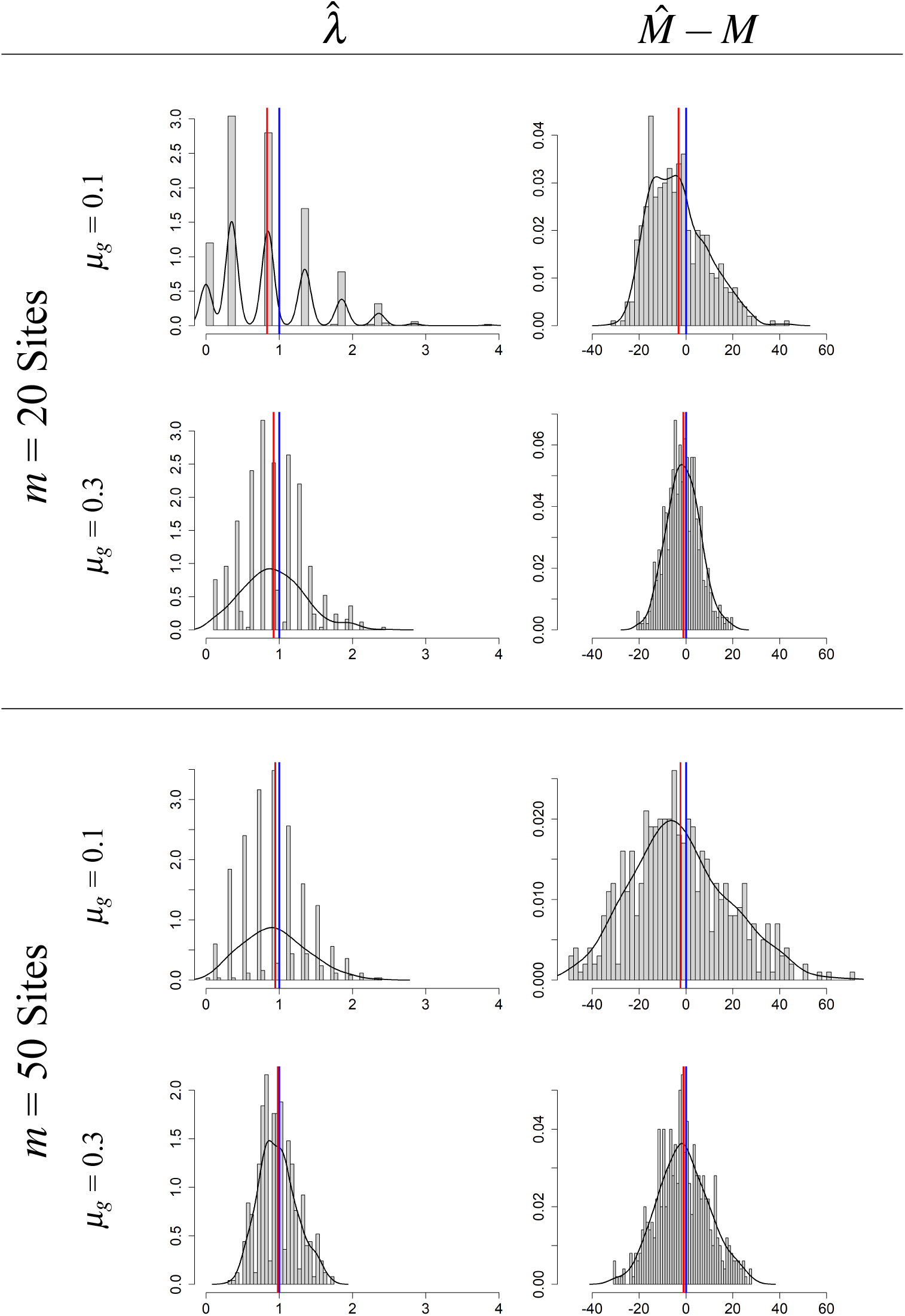
Histograms of estimated event rate (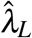 and 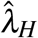) during simulations of a bimodal distribution for *λ*, and distribution of estimation errors for total number of events 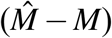. Vertical blue lines are the true parameter values. Vertical red lines mark the mean of simulation estimates. Black lines are a (Gaussian) kernel density estimate for the distribution.

Simulations that contained one low and one high rate (*λ*_*L*_ and *λ*_*H*_) also showed uni-modal distributions centered over the correct values of 1.0 and 10.0 (Figure 3). The distribution of the estimator for *λ*_*L*_ only overlapped that of the estimator for *λ*_*H*_ when *M* was low (20 sites) and *µ*_*g*_ = 0.1. The distribution of estimation errors surrounding the true value of *M* was unimodal and symmetric about zero in all cases despite the fact that true *M* was a mixture of Poisson distributions with means equal to 1 and 10. Bias of both 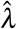 and 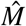 decreased as *M* or *µ*_*g*_ increased (Table 5). Estimated coverage of credible intervals of both *λ*_*H*_ and *M* was slightly above the nominal 90% level and varied from 93% to 96% (Table 5). Credible interval coverage for *λ*_*L*_ was only seriously low (62%) when the detection probably and number of sites were low. However, coverage trended toward the nominal level as detection probabilities increased.

**Figure 3:**
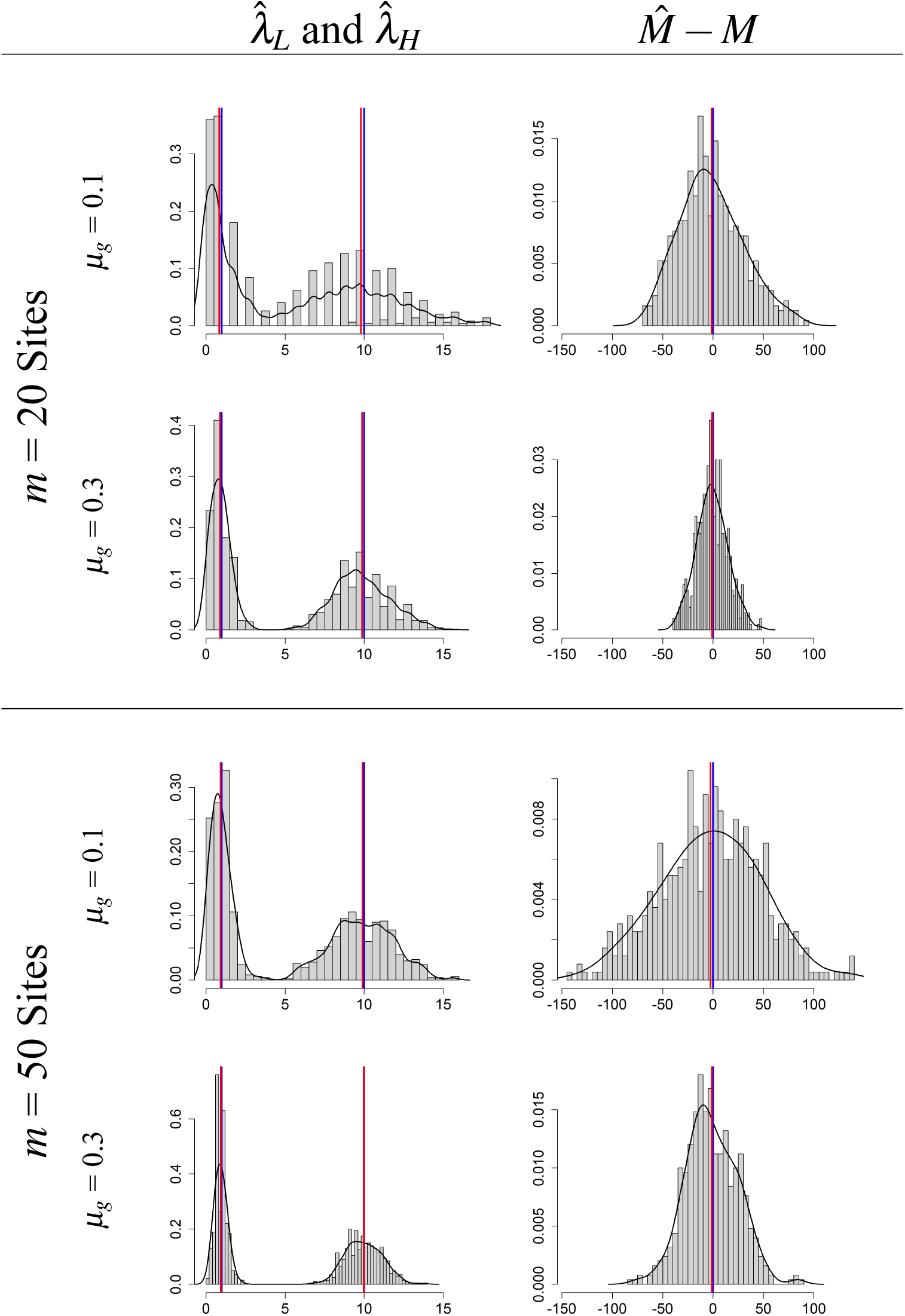
Histograms of estimated event rate (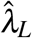 and 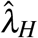) during simulations of a bimodal distribution for *λ*, and distribution of estimation errors for total number of events 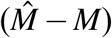. Vertical blue lines are the true parameter values. Vertical red lines mark the mean of simulation estimates. Black lines are a (Gaussian) kernel density estimate for the distribution.

Simulations containing two facilities with 40 and 10 turbines, respectively, and mean *g* values of 0.1 and 0.9, respectively, showed little bias and good confidence interval coverage when carcasses were consistently observed in a group. That is, the simulation showed little bias for the *λ* parameter associated with *g*_*i*_ = 0.9 (Simulation 3, Table 5) and confidence interval coverage for the group with higher detection rates was near the nominal level. But, bias was higher and confidence interval coverage exceeded the nominal level for the other (*g*_*i*_ = 0.1) group. The second group of the second run that contained only 10 turbines and *g*_*i*_ = 0.1 rarely observed carcasses and hence rarely contributed information to estimation of *λ*_*i*_. Still, percent bias for the rate parameter in this group was under 20% (Simulation 3, Table 5) illustrating the fact that EoAR produces accurate estimates when carcasses are observed. Due to the scarcity of carcasses in one group and lack of information to estimate *β*_*H*_, confidence interval coverage for total mortalities (*M*) suffered and exceeded the nominal level by 7% to 9%.

## 5 Discussion

The regression-type approach of EoAR provides increased flexibility relative to techniques that do not allow covariates, such as the Horvitz-Thompson (Huso, 2011) and straight EoA methods (Huso et al., 2015; Dalthorp, Huso, Dail, and Kenyon, 2014; Dalthorp, Huso, and Dail, 2017). Typically, analyses of wind-facility post-construction carcass counts estimate a mean rate and apply it to an entire facility or set of facilities. This approach is acceptable when all habitats and covariate combinations within or across facilities have been observed. When all turbines in a facility, or all turbines in all facilities, receive equal search effort, analysts can be reasonably comfortable that all covariate combinations were observed and that adjustment of a mean rate into the number of missed carcasses will yield accurate aggregate estimates. However, this mean-only approach is not adequate if important covariate combinations are not observed or if managers require estimates at facilities that are operated differently or that reside in substantially different environments. When carcass counts differ, or search effort is not uniform within facilities, or across facilities, extrapolation of a constant rate to additional sites can seriously bias overall mortality estimates. In addition, constant rates do not apply when future operations or habitat are known to change. Selecting an *a priori* probability sample of sites certainly diminishes the possibility of failing to observe important covariate combinations, but does not eliminate it. A better approach, and one that is consistent across multiple disciplines of ecology, is to identify factors causing variation in counts and to use those factors to improve the accuracy and precision of estimates. EoAR facilitates this approach by estimating direct relationships between covariate values and carcass counts.

EoAR is not the only analytical technique that estimates direct covariate relationships. At first glance, logistic and Poisson regression are viable alternative analysis techniques for wind facility monitoring data. If carcasses are common, logistic and Poisson regression can certainly be applied. But, the number of observed carcasses of many avian and bat species is relatively low and it is often unclear whether these models are seriously biased. That is, the level of carcass count required to produce unbiased logistic or Poisson regression models is unclear. Moreover, logistic regression cannot be applied when researchers find no carcasses (Huso, 2011). Assuming no complete or quasi-complete separation issues, weighted logistic regression (King and Zeng, 2001) can produce unbiased estimates of rare event probabilities, but only after difficult-to-assess exogenous information on the background proportion of successes in the population is incorporated into weights.

Like weighted logistic regression, EoAR also requires extra information in the form of detection probabilities (*g*) to complete estimation. However, unlike weighted logistic regression, EoAR’s extra information does not require abundance or a population-level proportion of success. EoAR only requires outside information on *g*, and *g* can usually be computed from the study design, searcher efficiency, and carcass removal trials.

Aside from being easy to compute because abundance is not required, the amount of information injected into an EoAR model by exogenous information on detection probabilities (*g*) is substantial and solves a number of problems associated with other techniques. First, outside information on detection makes a variable total (in this paper denoted *M*_*T*_) estimable in N-mixture models. Without an outside estimate of detection rate, information about variable totals and detection probabilities are confounded in the N-mixture model and cannot be separated (Barker et al., 2018). If true counts and detection probabilities are constant, an N-mixture model can estimate both *M*_*T*_ and *g* from a data set containing multiple observations because the first two sample moments are sufficient to estimate the two parameters of the binomial distribution.

Second, information provided by detection probabilities allows an EoAR model to be fitted when the data are completely or quasi-completely separated among covariate combinations. With information on detection, zero percent success in one or more covariate combinations is a valid observation and it can be adjusted correctly. EoAR is then a viable method for resolving complete and quasi-complete separation problems in binomial models if detection probabilities (i.e., *g*) are available from some outside source. A couple of methods for resolving complete or quasi-complete separation that do not require detection probabilities are reviewed in Heinze (2006).

Detection probability *g* is, in concept, similar to the detection probability parameters in other ecological field sampling techniques like distance sampling (Borchers, Zucchini, and Fewster, 1998; Buckland, Rexstad, et al., 2015; Buckland, Anderson, et al., 2004), occupancy analyses (MacKenzie and Kendall, 2002; MacKenzie, Nichols, Lachman, et al., 2002; MacKenzie, Nichols, Sutton, et al., 2005), and capture-recapture methods (Amstrup, McDonald, and Manly, 2005; Borchers and Efford, 2008; Efford, Borchers, and Byrom, 2009). Those methods, and EoAR, require probability of detection conditional on availability (i.e., P(*detect*|*available*)) to adjust observed counts for missed targets. The difference between distance, occupancy, and capture-recapture analysis and EoAR is that the former techniques estimate detection probabilities from the same data available for abundance estimation. Indeed, one could argue that the key feature of these non-EoAR techniques is that they estimate probability of detection from a single data set so that observed counts can be adjusted into density, occupancy, or abundance. EoAR uses study design elements such as search timing, search frequency, and the proportion of carcasses sampled, along with measured quantities like average carcass lifetime and searcher efficiency, to estimate probability of detection.

If components of *g* are unknown (e.g., carcass persistence rates), computation of EoAR’s detection probability can be difficult. EoAR requires at least that detection probabilities be bracketed by lows and highs prior to analysis. If low and high values are identified, the variation in *g* between the lows and high can be built into *g*’s Beta distribution by setting *α* and *Ψ* accordingly. If absolutely no information is available on the range of detection probabilities, it is tempting to hypothesize a uniform distribution for *g* and set both *α* and *Ψ* to 1.0. However, in this special case no additional information is injected into the model and the convergence of *λ* (and *M*_*T*_) depends solely on variation in the observed counts across covariate combinations. In the case when absolutely no information on *g* is available, practitioners should probably reject our model in favor of standard logistic or Poisson regression and pool counts until *C*_*i*_ > 0 for all covariate combinations.

Example 2 showed reasonable results when only one carcass at one turbine among 811 was observed. Example 2 also illustrated that *λ*’s prior can have a major impact on EoAR’s final results, especially when carcass counts are low. In general, we have found that EoAR’s results are particularly sensitive to *λ*’s prior when carcass counts are exactly zero. When little or no information is collected (i.e., no carcasses) the only information EoAR has is *λ*’s prior. If *λ*’s prior is vague and *C* = 0, EoAR will quite reasonably estimate extremely small rates and essentially no mortalities. In contrast, EoA software currently always uses a distribution proportional to the Jefferey’s prior for *λ* (Dalthorp, Huso, and Dail, 2017; Dalthorp, 2019). EoA’s prior always places a significant amount of weight on zero, but also places a significant amount of weight on positive *λ*’s. EoA’s Jeffery’s prior is weakly informative in the sense that it is much less vague than EoAR’s vague priors. Using this weakly informative prior, EoA produces positive (non-zero) upper bounds for fatalities when *C* = 0 that depend heavily on *g*. We believe positive upper bounds on mortality when *C* = 0 are acceptable provided practitioners understand the situation and agree with the weakly informative prior on *λ*. We plan to implement a weakly informative prior in the EoAR package for the case when *C* = 0, but it is more difficult for EoAR due to the linear model (i.e., priors on the *β* coefficients need to be weakly informative). More generally, we caution practitioners to always carefully consider *λ*’s prior, and particularly when no carcasses are found.

In addition to its other features, EoAR is useful for planning and study design. Studies can generally control probability of detection, at least absent budgetary constraints, and this allows planners to limit or control uncertainty in the form of credible interval widths. Studies that can accept large amounts of uncertainty can design data collection efforts that result in low *g* values because low *g* values generally produce wide credible intervals. Studies that require high precision surrounding the estimated total number of fatalities must allocate effort wisely and design data collection efforts to achieve high *g* values. At a minimum, study designers can hypothesize various *g* values, compute credible interval widths and budgetary requirements associated with each, and weigh the associated field costs against precision of the final estimates.

In summary, EoAR provides viable estimates and upper-bounds for the number of fatalities in all situations assuming appropriate prior distributions are used. EoAR has the desirable feature of relating fatality rates to study covariates, which in turn increases precision. EoAR models are robust to quasi-complete separation issues and are relatively easy to compute using the R package referenced in Appendix S1. The only drawback of EoAR is that separate estimates of detection probabilities are required, and for wind facility monitoring these require additional field trials to establish searcher efficiency and carcass removal. This additional information on detection probabilities is worth acquiring because doing so generally requires less effort than acquiring separate information required by alternative techniques.

## 6 Acknowledgements

The authors wish to thank Kraig McPeek, U.S. Fish and Wildlife Service, for his encouragement and insightful comments. We also wish to thank the dozens of Western EcoSystems Technology and MidAmerican field personnel who diligently collected the field data used in the examples. Two anonymous referees provided extensive comment on the draft manuscript which greatly improved the final product.

## References

1. Albert, A. and J. A. Anderson (1984). “On the existence of maximum likelihood estimates in logistic regression models”. Biometrika 71 (1), pp. 1–10.

2. Amstrup, S. C., T. L. McDonald, and B. F. J. Manly (2005). Handbook of Capture-Recapture Analysis. Princeton, NJ: Princeton University Press, p. 313.

3. Barker, R. J., M. R. Schofield, W. A. Link, and J. R. Sauer (2018). “On the reliability of N-mixture models for count data”. Biometrics 74 (1), pp. 369–377.

4. Borchers, D. L. and M. G. Efford (2008). “Spatially Explicit Maximum Likelihood Methods for Capture-Recapture Studies”. Biometrics 64 (2), pp. 377–385.

5. Borchers, D. L., W. Zucchini, and R. M. Fewster (1998). “Mark-Recapture Models for Line Transect Surveys”. Biometrics 54 (4), pp. 1207–1220.

6. Buckland, S. T., E. A. Rexstad, T. A. Marques, and C. S. Oedekoven (2015). Distance Sampling: Methods and Applications, pp. 1–277.

7. Buckland, S. T., D. R. Anderson, K. P. Burnham, J. L. Laake, D. L. Borchers, and L. Thomas (2004). Advanced distance sampling. Oxford University Press.

8. Dalthorp, D., L. Madsen, M. Huso, P. Rabie, R. Wolpert, J. Studyvin, J. Simonis, and J. Mintz (2018). GenEst statistical models-A generalized estimator of mortality: U.S. Geological Survey Techniques and Methods. Tech. rep. A2. U.S Geological Survey.

9. Dalthorp, D. (2019). eoa: Wildlife mortality estimator for scenarios with low fatality rates and imperfect detection. R package version 2.0.7.

10. Dalthorp, D. and M. Huso (2015). A Framework for Decision Points to Trigger Adaptive Management Actions in Long-Term Incidental Take Permits. Tech. rep. January, p. 88.

11. Dalthorp, D., M. Huso, and D. Dail (2017). Evidence of absence (v2.0) software and user guide: U.S. Geological Survey Data Series 1055. Tech. rep. U.S Geological Survey.

12. Dalthorp, D., M. Huso, D. Dail, and J. Kenyon (2014). Evidence of absence software and user guide: U.S. Geological Survey Data Series 881. Tech. rep. U.S Geological Survey.

13. Dalthorp, D., J. Simonis, L. Madsen, M. Huso, P. Rabie, J. Mintz, R. Wolpert, J. Studyvin, and F. Korner-Nievergelt (2020). GenEst: Generalized Mortality Estimator. R package version 1.4.4.

14. Efford, M. G., D. L. Borchers, and A. E. Byrom (2009). “Density Estimation by Spatially Explicit Capture-Recapture: Likelihood-Based Methods”. In: Modeling Demographic Processes In Marked Populations. Ed. by D. Thomson, E. Cooch, and M. Conroy. Vol. 3. Environmental and Ecological Statistics. Springer US, pp. 255–269.

15. Erickson, W. P., M. M. Wolfe, K. J. Bay, D. H. Johnson, and J. L. Gehring (2014). “A comprehensive analysis of small-passerine fatalities from collision with turbines at wind energy facilities”. PLoS ONE 9 (9).

16. Gelman, A., J. B. Carlin, H. S. Stern, and D. B. Rubin (2004). Bayesian Data Analysis: Second Edition. CRC Press.

17. Heinze, G. (2006). “A comparative investigation of methods for logistic regression with separated or nearly separated data”. Statistics in Medicine 25 (24), pp. 4216–4226.

18. Horvitz, D. G. and D. J. Thompson (1952). “A Generalization of Sampling Without Replacement From a Finite Universe”. Journal of the American Statistical Association 47 (260), p. 663.

19. Huso, M. M. P. (2011). “An estimator of wildlife fatality from observed carcasses”. Environmetrics 22 (3), pp. 318–329.

20. Huso, M. M. P., D. Dalthorp, D. Dail, and L. Madsen (2015). “Estimating wind-turbine-caused bird and bat fatality when zero carcasses are observed”. Ecological Applications 25 (5), pp. 1213–1225.

21. Jain, A. (2005). “Bird and Bat Behavior and Mortality at a Northern Iowa Windfarm”. PhD thesis. Iowa State University, Ames Iowa.

22. Kéry, M. (2018). “Identifiability in {N-mixture} models: a large-scale screening test with bird data”. Ecology 99 (2), pp. 281–288.

23. Kéry, M. and J. A. Royle (2016). Applied Hierarchical Modeling in Ecology. Vol. 1. San Diego, California: Academic Press, p. 735.

24. King, G. and L. Zeng (2001). “Logistic Regression in Rare Events Data”. Political analysis 9 (2), pp. 137–163.

25. Korner-Nievergelt, F., P. Korner, O. Behr, I. Niermann, R. Brinkmann, and B. Hellriegel (2011). “A new method to determine bird and bat fatality at wind energy turbines from carcass searches.” Wildlife Biology 17 (4), pp. 350–363.

26. Korner-Nievergelt, F., T. Roth, S. Von Felten, J. Guélat, B. Almasi, and P. Korner-Nievergelt (2015). Bayesian data analysis in ecology using linear models with R, BUGS, and Stan. Academic Press.

27. Kruschke, J. K. (2011). Doing Bayesian data analysis : a tutorial with R and BUGS. Academic Press, p. 653.

28. Lemoine, N. P. (2019). “Moving beyond noninformative priors: why and how to choose weakly informative priors in bayesian analyses”. Oikos 128 (7), pp. 912–928.

29. Lohr, S. L. (2010). Sampling: Design and Analysis. Second. Brooks/Cole.

30. Maalouf, M. and T. B. Trafalis (2011). “Robust weighted kernel logistic regression in imbalanced and rare events data”. Computational Statistics and Data Analysis 55 (1), pp. 168–183.

31. MacKenzie, D. I. and W. L. Kendall (2002). “How should detection probability be incorporated into estimates of relative abundance”. Ecology 83 (9), pp. 2387–2393.

32. MacKenzie, D. I., J. D. Nichols, J. E. Hines, M. G. Knutson, and A. B. Franklin (2003). “Estimating Site Occupancy, Colonization, and Local Extinction When a Species is Detected Imperfectly”. Ecology 84 (8), pp. 2200–2207.

33. MacKenzie, D. I., J. D. Nichols, G. B. Lachman, S. Droege, J. A. Royle, and C. A. Langtimm (2002). “Estimating site occupancy rates when detection probabilities are less than one”. Ecology 83 (8), pp. 2248–2255.

34. MacKenzie, D. I., J. D. Nichols, N. Sutton, K. Kawanishi, and L. L. Bailey (2005). “Improving Inferences in Population Studies of Rare Species That Are Detected Imperfectly”. Ecology 86 (5), pp. 1101–1113.

35. Maratea, A., A. Petrosino, and M. Manzo (2014). “Adjusted {F-measure} and kernel scaling for imbalanced data learning”. Information Sciences 257, pp. 331–341.

36. McCullagh, P. and J. Nelder (1989). Generalized Linear Models. 2nd. London: Chapman and Hall.

37. Pavlacky, D. C., J. A. Blakesley, G. C. White, D. J. Hanni, and P. M. Lukacs (2012). “Hierarchical multi-scale occupancy estimation for monitoring wildlife populations”. Journal of Wildlife Management 76 (1), pp. 154–162.

38. Plummer, M. (2003). “JAGS: A program for analysis of Bayesian graphical models using Gibbs sampling”. Proceedings of the 3rd International Workshop on Distributed Statistical Computing (DSC 2003), pp. 20–22.

39. Reyes, G., M. Rodriguez, K. Lindke, K. Ayres, M. Halterman, B. Boroski, and D. Johnston (2016). “Searcher Efficiency and Survey Coverage Affect Precision of Fatality Estimates”. The Journal of Wildlife Management 80 (8), p. 1488.

40. Särndal, C.-E., B. Swensson, and J. Wretman (1992). Model Assisted Survey Sampling. Springer Series in Statistics. New York: Springer-Verlag.

41. Shoenfeld, P. (2004). Suggestions Regarding Avian Mortality Extrapolation. Tech. rep. FPL Energy.

42. West Virginia Highlands Conservancy, HC70, Box 553, Davis, West Virginia, 26260.

43. Silverman, B. W. (1998). Density Estimation for Statistics and Data Analysis. New York: Routledge.

44. Simpson, D., H. Rue, A. Riebler, T. G. Martins, and S. H. Sørbye (2017). “Penalising model component complexity: a principled, practical approach to constructing priors”. Statistical Science 32 (1), pp. 1–28.

45. Spiegelhalter, D. J., N. G. Best, B. P. Carlin, and A. van der Linde (2014). “The deviance information criterion: 12 years on (with discussion)”. Journal of the Royal Statistical Society, Series B 76 (3), pp. 485–493.

46. Studyvin, J., P. Rabie, and D. Riser-Espinoza (2020). windAC: Area Correction Methods. R package version 1.2.2.

