## Appendix S1 for "Evidence of Absence Regression: A Binomial N-Mixture Model for Estimating Fatalities at Wind Energy Facilities"

### Evidence of Absence Regression: A Binomial N-Mixture Model for Estimating Fatalities at Wind Energy Facilities

Trent McDonald[1], Kimberly Bay[2], Jared Studyvin*[2], Jesse Leckband[3], Amber Schorg[4], and Jennifer McIvor[5]

[1]Western Ecosystems Technology, Inc., Cheyenne, WY, USA and McDonald Data Sciences, LLC, Laramie, WY, USA
[2]Western Ecosystems Technology, Inc., Cheyenne, WY, USA
[3]MidAmerican Energy Company, Des Moines, IA, USA
[4]Rock Island Field Office, U.S. Fish and Wildlife Service, Moline, IL, USA
[5]Berkshire Hathaway Energy Company, Des Moines, IA, USA,

2 February 2021

Journal: Ecological Applications

### S1 EoAR's Directed Acyclic Graph

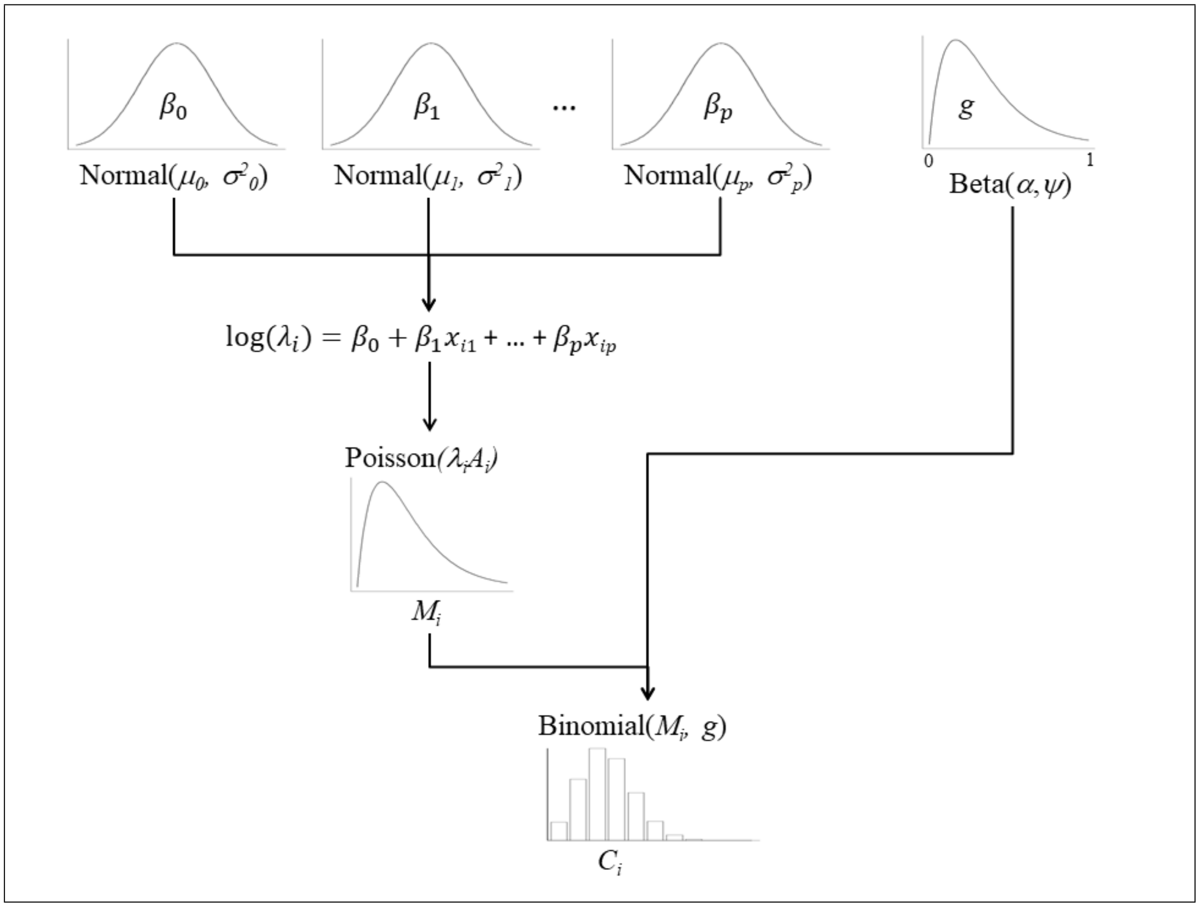


Figure **S1:** A directed acyclic graph of the Evidence of Absence Regression (EoAR) model loosely based on the directed acyclic graphs of Kruschke (2011). Top row of distributions represent the priors and can be either vague or informed. Data are carcass counts $C_i$. Scaling offsets are $A_i$. Beta parameters $\alpha$ and $\psi$ must be estimated from the study design (e.g., with routines in R package `GenEst`).

### S2 EoAR Code

R code to estimate an EoAR model is available from the author's GitHub site: `https://github.com/tmcd82070/EoAR`.

Code for the simulations and subsequent analysis, as well as the Iowa bat carcass data set, can be obtained from the author's Dryad data repository:

`https://doi.org/10.5061/dryad.2rbnzs7jh`
or
`https://datadryad.org/stash/share/r0UoSyX2ZB9UuovHz4mOHSk97TGBXiHegBUGzttouqg`
